# DarkQ: Continuous genomic monitoring using message queues

**DOI:** 10.1101/2020.11.12.379560

**Authors:** A. Viehweger, C. Brandt, M. Hölzer

## Abstract

**Motivation:** Newly sequenced genomes are often not noticed by potential stakeholders because submission to public databases is delayed, and search options are limited. However, the discovery of genomes can be vital: In pathogen outbreaks, fast updates are essential to coordinate containment efforts and prevent spread.

**Results:** Here we introduce DarkQ, a message queue that allows for instant sharing and discovery of genomes.

**Availability:** DarkQ is released under the BSD-2 license at github.com/phiweger/darkq.

## Introduction

Many bioinformatic tasks complement newly sequenced genomes with existing, publicly available ones. For example, when reconstructing a local pathogen outbreak, screening similar genomes can discover related ones from other sampling sites, such as hospitals nearby, and can significantly affect public health response.^1,2^ However, no tool exists to monitor newly sequenced genomes and automatically identify those of interest to the user. A long delay until they are publicly available explains why many outbreak studies are retrospective and offer limited practical value to the associated outbreak response. Several components are needed: First, a “publisher” needs to be able to send genomic “messages” to a “consumer” using a simple and secure interface. Second, the genomic “message” may require limited space to avoid upload problems or extensive storage infrastructure. Third, a mechanism is needed to route messages only to interested parties, e.g., consumers that search for genomes of a particular species in a specific geography. Lastly, on receiving a relevant message, download of the associated genome should be possible. Several projects currently develop ways to share genomes effectively (github.com/dib-lab/wort, github.com/will-rowe/stark). However, to our knowledge, ours is the first end-to-end solution available to users.

## Implementation

DarkQ is implemented using the Nextflow workflow manager to ensure a robust, reproducible, and portable application.^3^ The user interface of DarkQ is similar to the popular file system service “Dropbox”: The content of a “send” directory is tracked. When a genome is added to it, it is first compressed (“sketched”) using the MinHash algorithm^4,5^ (sourmash, v3.5). The reduction in file size by orders of magnitude allows for efficient transmission. Together with metadata and inferred taxonomy (using sourmash), the genome sketch constitutes a “message” (Supplement, Fig. S1, A). The receiving message queue then uses the Advanced Message Queuing Protocol (AMQP)^6^ to route messages (implementation: github.com/rabbitmq, v3.8.9) onto queues, i.e., sequential groups of messages. The original genome is uploaded (“pinned”) to a decentralized, peer-to-peer network (IPFS, v0.7).^7^ Its content-based address is part of the genome message.

The consumer can subscribe to messages using an arbitrary number of filters, so-called “routing keys”. Each routing key is unique and has five properties: name of sender (e.g. “phiweger”), country code (e.g. “DE”), taxon status (“found” or “mystery”), taxon level (either one of superkingdom, phylum, class, order, family, genus, species, strain) and taxon name at that level (e.g. “Klebsiella” for genus) – these are adapted from and must conform to the Genome Taxonomy Database (GTDB, release 89).^8^ For example, “phiweger.DE.*.genus.klebsiella” would select all isolate genomes of the stated genus from Germany sent by the author.

Because we can estimate genome similarity using MinHash sketches,^4^ the consumer can quickly filter the received genome messages using target genomes, e.g., those belonging to a local pathogen outbreak or current research project. If this filter is passed, then the genome is automatically downloaded from the peer-to-peer network using its content hash address, which at the same time locates and validates the downloaded file. If multiple users pin the genome, download speed can increase substantially. A downstream workflow can then be connected to refine these genomes’ analysis further, enabling a complete monitoring system.

## Usage scenario

To test DarkQ in a monitoring system, we collected and sent onto DarkQ 9,415 genomes of *Klebsiella pneumoniae*; a pathogen considered an urgent global threat due to extensive antimicrobial drug resistance (CDC, AR threats report, 2019).^9^ We simulated a consumer subscribing to all messages from the Klebsiella genus and filtering the received messages using an isolate from a local outbreak at a large tertiary hospital in 2010.^10^ 1,461 messages met both routing key and minimum genome similarity criteria of 0.97 at a k-mer size of 51, typically used to estimate the genomic distance at the strain level.^11^ After downloading the original genomes from the peer-to-peer network, they were further filtered and refined, resulting in a time-dated phylogeny (Fig. S1, B and methods in supplement). The consumer thus received genomes from a total of 26 studies. Two of these studies contained genomes that belonged to the same outbreak clone the consumer used to filter the genomes. Further work is needed to investigate this relationship more thoroughly, however, an initial assessment was already possible by utilizing the mechanics implemented in DarkQ

## Conclusion

DarkQ allows to monitor genomic data with a simple user interface, efficient genome compression, filter-based message routing, and fast download of corresponding genomes using a decentralized peer-to-peer network. The proof-of-concept outlined here scales to thousands of genomes and could be particularly valuable in the context of pathogen outbreaks. However, our approach can be used to disseminate research more broadly.

## Acknowledgements

We thank Luiz Irber and C. Titus Brown (University of California, Davis) for insightful discussions of the concepts discussed in this article.

## Supplement

**Fig. S1:**
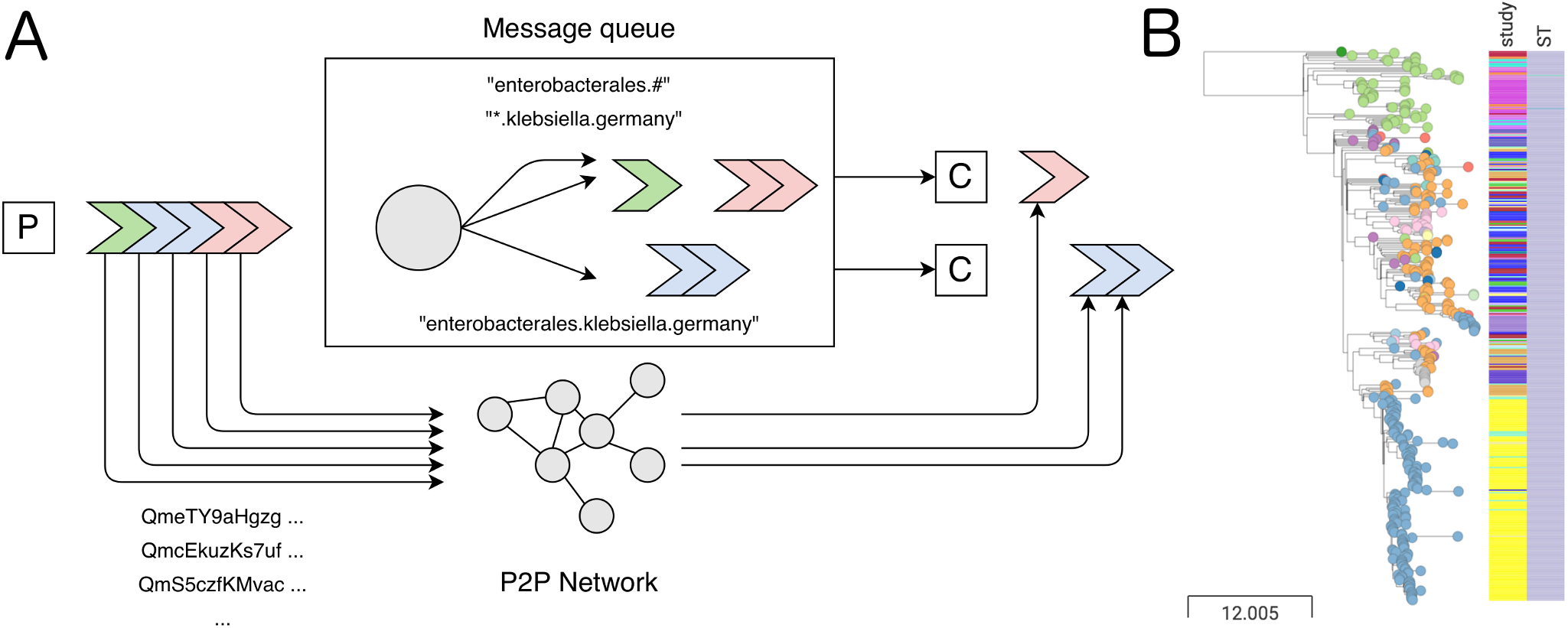
**(A)** Architecture of DarkQ: Compressed genome “messages” (colored arrows) are sent by a publisher (P) onto the message queue. A router (circle) distributes the messages to queues via routing keys (annotated arrows). Consumers (C) can use these keys to receive only a subset of messages and then further filter them with target genomes using MinHash sketches. In parallel, the genomes from the publisher are uploaded on a decentralized peer-to-peer (P2P) network. Once messages pass through the consumer’s filters, they are automatically downloaded from the P2P network. This architecture allows the effective distribution of newly sequenced genomes and enables continuous monitoring, e.g., in outbreak scenarios. **(B)** Use case simulation: A hospital becomes aware of a local outbreak of an XDR *Klebsiella pneumoniae* (Kp) isolate of subtype (ST, right metadata column) 258 carrying a plasmid-encoded KPC-2 carbapenemase. Using DarkQ, we identified 431 genomes from several countries (leaf colors) from 26 studies (left metadata column) with an average nucleotide identity (ANI) > 99.98% and identical resistance and capsule patterns (not shown). A time-dated phylogeny revealed several non-local isolates, suggesting that the outbreak reached further than previously assumed. An interactive version of the data can be found at microreact.org/project/facEFbDrgwgp9aX97nvpHq. Scale in number of SNVs.

## Supplementary Methods

Exact nucleotide identity was calculated using FastANI v1.31 (1). Single nucleotide variants (SNVs) were called using Snippy v4.6 (2). Intervals of increased SNV density likely due to recombination were masked using Gubbins v2.4.1 (3), and the remaining SNVs were parsed using SNP-sites v2.4 (4). A maximum-likelihood tree was built using FastTree v2.1.10 (5). A time-dated phylogeny was modeled using Treetime 0.7.6 (6). For interactive visualization, we used the Microreact web service (7).

## Notes

https://github.com/phiweger/darkq

### Competing Interest Statement

The authors have declared no competing interest.

### Summary of Updates

Abstract changed

